# LIMK Inhibition and Metformin Block Mitochondrial Transfer Overcoming Macrophage Driven Therapy Resistance in Acute Myeloid Leukaemia

**DOI:** 10.64898/2026.02.03.702377

**Authors:** Ebubechukwu Nwarunma, Katerina E. Miari, Athanasia Papadopoulou, Stefan Corradini, Glen Watt, Samantha Hurwitz, Tatiana Fourfouris, Ki Jun Lee, Xenia Bubnova, Rebecca Briggs, Carl S Goodyear, Theodoros Simakou, Marcus Doohan, Lucy MacDonald, Mariola S. Kurowska-Stolarska, Timothy J. Humpton, Victoria L. Campbell, Lesley M. Forrester, Ken I. Mills, Katrina Lappin, Valerie A. Ferro, Yong-Mi Kim, Helen Wheadon, Mark T.S. Williams

**Affiliations:** Research Centre for Health, School of Health and Life Sciences, Department of Biological and Biomedical Sciences, Glasgow Caledonian University, Cowcaddens Road, Glasgow, G4 0BA, UK; School of Cancer Sciences, College of Medical, Veterinary and Life Sciences, University of Glasgow, Glasgow, UK; Children’s Hospital Los Angeles, Keck School of Medicine, University of Southern California, 4650 Sunset Boulevard. Los Angeles, CA 900327; Johnson Cancer Research Centre, Queen’s University Belfast; School of Infection and Immunity, College of Medical, Veterinary and Life Sciences, University of Glasgow, Glasgow, UK; Cancer Research UK Scotland Institute, Glasgow, UK; Western General Hospital, Edinburgh, United Kingdom; Centre for Regenerative Medicine, Institute of Regeneration and Repair, The University of Edinburgh, Edinburgh, Scotland, United Kingdom; Strathclyde Institute of Pharmacy and Biomedical Sciences, University of Strathclyde, Glasgow, U.K

**Keywords:** AML, macrophages, chemoresistance, mitochondrial transfer, ROS

## Abstract

Chemoresistance is a major contributor to poor clinical outcomes in AML patients and can arise from interactions between AML cells and the bone marrow microenvironment (BME). How immune cells, particularly macrophages (Mφs), facilitate this process requires better clarification. This study shows that M2-like Mφs protect AML cells from apoptosis induced by daunorubicin (DNR) and cytarabine (Ara-C). This protection occurs via co-culture and is linked to enhanced mitochondrial transfer from Mφs to AML cells. Mφs interacted with AML cells via tunneling nanotube (TNT)-like structures. Furthermore, inhibition of mitochondrial transfer using cytochalasin B reduced the protective effect, indicating that mitochondria mediate this process. Mφs transferred functional mitochondria to AML cells as evidenced by enhanced metabolic capacity and reduced reactive oxygen species levels in AML cells under chemotherapy stress. TH-257 (LIMK inhibitor) and metformin blocked mitochondrial transfer and Mφ-driven chemoprotection. Moreover, increased transcript expression levels of RhoC and cofilin correlate with inferior overall survival in AML patients. These findings suggest that M2-like Mφs contribute to chemoresistance through TNT-mediated mitochondrial transfer and the LIMK-Cofilin pathway, identifying potential therapeutic targets to circumvent chemoresistance in AML.

## INTRODUCTION

Acute Myeloid Leukemia (AML) is a genetically diverse, multi-clonal malignancy characterized by the accumulation of immature myeloblasts in the bone marrow (BM) and peripheral blood. Although targeted therapies have emerged, traditional chemotherapy with daunorubicin (DNR) and cytarabine (Ara-C) remains the foundation of AML induction treatment. However, 50% to 70% of adult AML patients who achieve a complete remission (CR) after induction therapy will eventually relapse due to the development of therapy-resistant leukemic stem and progenitor cells. Within the bone marrow microenvironment (BME), AML cells interact with resident cells such as mesenchymal stromal cells (MSCs) and immune cells, including macrophages (Mφs) (1–3). AML blasts reprogram the BME to support leukaemia progression at the expense of normal haematopoiesis. These stromal interactions impair response to both chemotherapy and targeted agents, contributing to treatment failure (4, 5). Macrophages, highly plastic immune cells, can polarize into M1-like (pro-inflammatory, CD86^+^) or M2-like (anti-inflammatory, CD163^+^CD206^+^) phenotypes. AML patient BM shows elevated levels of M2-like Mφs, which are associated with poor responses to induction therapy (6, 7). An M2-like macrophage gene signature including *CD163*, *CD206*, *FGR*, *CD52*, *RASA3*, and *GSK1B* has been linked to reduced overall survival (OS) in AML (8). AML blasts secrete factors like macrophage migration inhibitory factor (MIF) that reprogram Mφs toward the M2 phenotype, which in turn reduces AML cell sensitivity to targeted therapy (9, 10). Recent studies have also shown that mitochondria are transferred from stromal cells-such as fibroblasts (11), MSCs (12), and macrophages-to AML cells (8).

This transfer can enhance AML cell survival during Ara-C therapy (11). CD38 is implicated in mediating mitochondrial transfer from MSCs (12). Despite these insights, the role of M2-like CD163^+^CD206^+^ macrophages in driving chemoresistance to DNR and Ara-C through direct contact and mitochondrial transfer remains unclear. In this study, we evaluated the impact of M2-like Mφs on AML therapy response. Using an *ex vivo* co-culture system with AML cells, human monocyte-derived Mφs (MDMs), iPSC-derived Mφs, and murine BMDMs, we showed M2-like Mφs significantly protect AML cells from DNR-induced apoptosis. Mechanistically, this likely occurs via tunneling nanotube (TNT)-mediated mitochondrial transfer, reducing reactive oxygen species (ROS) in AML cells. An inhibitor of LIMK1/2 (TH-257), and metformin suppressed mitochondrial transfer and chemoprotection. Our findings reveal that targeting TNT-mediated mitochondrial transfer may offer a novel strategy to overcome chemoresistance in AML.

## METHODS

### Cell lines and Primary cells

Human AML cell lines U937, THP-1, and KG1a (ATCC) were cultured in complete RPMI 1640 (cRPMI, Gibco) with 10% v/v FBS (20% for KG1a), 1% v/v penicillin/streptomycin, and 1% v/v L-glutamine (Invitrogen). Primary AML cells or BM aspirates were obtained from apheresis products of patients that provided informed consent and IRB approval from the Northern Ireland Biobank and the NHS Greater Glasgow and Clyde Biorepository. See **Supplementary Table 1** for clinical information of primary AML specimens. Primary human AML cells were stored in Cryostor CS10 (Stemcell Technologies) and then cryo-preserved in liquid nitrogen. Freshly thawed cells were cultured in Serum Free Media (SFM) at 37°C, in 5% CO_2_ incubator. SFM is composed of IMDM (Gibco), 20% BIT-9500 (Stemcell Technologies), 10 mg/ml LDL (Low Density Lipoprotein, Millipore), 55 mM 2-Mercaptoethanol (GIBCO) and 1% v/v Pen/Strep (Gibco).

### Generation of human CD14^+^ monocyte derived macrophages

Peripheral blood was obtained from healthy donors (Scottish Blood Transfusion Service). CD14⁺ monocytes were isolated using microbeads (Miltenyi Biotec) and cultured at 1x10^6^ cells/mL in cRPMI with 100 ng/mL M-CSF (PeproTech) for 7 days. Media was replenished, and cells further cultured for 3 days to generate M2-like macrophages (CD206⁺).

### Confirming M2-like phenotype of MDMs

Day 10 MDMs were assessed via FACS for M2 (CD206) and M1 (CD86) markers. Macrophage conditioned media (Mφ-CM) and cRPMI were analyzed via a 30-Plex cytokine/chemokine panel (Thermo Fisher) on a Luminex LX-200 instrument.

### Flow cytometry

Apoptosis was measured using Annexin V and Fixable Viability Dye eFluor™ 450 (eBioscience) or DAPI. ROS levels were assessed using 7.5 μM CellROX Deep Red reagent (Thermo Fisher). Data were acquired on a MACSQuant® Analyzer 10 (Miltenyi Biotec) with 20,000 gated events and analyzed using FlowJo v10.5.3 (Tree Star Inc). Unless noted, results are reported as median fluorescence intensity (MFI).

### Macrophage Impact on Chemotherapy-Induced Apoptosis

AML cells were cultured in cRPMI, 50% Mφ-CM, or co-cultured 1:1 with Mφs ± daunorubicin (DNR) for 24-72h. Apoptosis was assessed as stated above. AML lines were gated on CD14-.

### Mitochondrial Transfer Assessment

Day 9 M2-like Mφs were washed twice, stained with MitoTracker Deep Red (MTDR) FM (1:1000) for 30 min, washed twice again, and incubated for 24h-72h. CFSE-labelled AML cells were co-cultured with Mφs, and mitochondrial transfer was assessed by detecting MitoTracker signal in CFSE⁺ AML cells.

### Tandem Mass Tag Mass Spectroscopy (TMT-MS)

CFSE-labelled AML cell lines were co-cultured directly with M2-like Mφs for 24-72h. CD14- AML cells were then isolated from CD14+ Mφs, via the use of CD14 microbeads (Miltenyi Biotec), with a purity of >90%. Proteomic profiling was then conducted on purified AML whole cell lysates (WCLs), via a TMT11plex, in collaboration with Dr Richard Burchmore (Glasgow Polyomics, University of Glasgow)(13).

### Generation of macrophage CM

Mφ-CM was collected on day 10, centrifuged (2000 rpm, 5 min), filtered through 0.22 µm filters (Millipore), and stored at 4°C for up to 24h.

### Western blotting

Cell lysis was performed using RIPA buffer with protease and phosphatase inhibitors (Sigma). Lysates were sonicated (3x1 sec bursts), and total protein measured by BCA assay (Thermo Fisher). Mφ-CM or 25 µg of total protein was resolved on 4-12% bis-tris gels (Invitrogen), transferred using iBlot, and probed with 1:1000 diluted antibodies (Cell Signaling Technologies); Phospho-Cofilin (Ser3) (77G2) Rabbit Monoclonal Antibody #3313, Cofilin (D3F9) Rabbit Monoclonal Antibody #5175 and Anti-rabbit IgG, HRP-linked Antibody #7074. Blots were visualized using West FEMTO supersignal (Thermo Fisher) and LI-COR Odyssey Fc imaging system.

### Generation of iPSC-derived macrophages

Human iPSC line SFCi55 (Gift from Dr. Luca Cassetta) were expanded in Stemflex medium (Thermofisher) and used to generate embryoid bodies and iMacs precursors using established protocols (14). iMac were differentiated from precursors for 7 days using 100 ng/mL M-CSF, and evaluated by flow cytometry (FACSCanto, BD Biosciences) before maturation.

### Murine Bone Marrow-Derived Macrophages (BMDMs)

BMDMs were generated from BM harvested from tibias and femurs of C57BL/6 mice following published methods (15).

### Mitochondrial Respiration

AML cells, following 24h co-culture with M2-like MDMs, were plated in Cell-Tak-coated wells of a 96-well plate and analyzed using the Seahorse XFp Analyzer and XF Mito Stress Test Kit (Agilent) as previously published (16). Media was supplemented with glucose (11 mM), L-glutamine (2 mM), and sodium pyruvate (1 mM). Mitochondrial function was assessed by sequential injection of oligomycin (1.5 µM), FCCP (1 µM), and a mix of antimycin and rotenone (1 µM). Data were normalized to cell counts using Wave software, with five technical replicates performed per condition in each independent experiment.

### Murine mtDNA/nDNA Determination

Total genomic DNA was purified via the PureLink Genomic DNA Mini Kit (Invitrogen). It was then quantified, diluted to 1 ng/mL and used to determine the relative copy number of the murine mitochondrial DNA (COX II) and nuclear DNA [Amyloid Precursor Protein (APP)] by Quantitative PCR. The samples were run on a QuantStudio 5 Real-Time PCR system (Thermo Fisher Scientific) using TaqMan FAST advanced master mix and the primer/probe combinations listed in **Table 1**, as previously published (16).

**Table 1.**
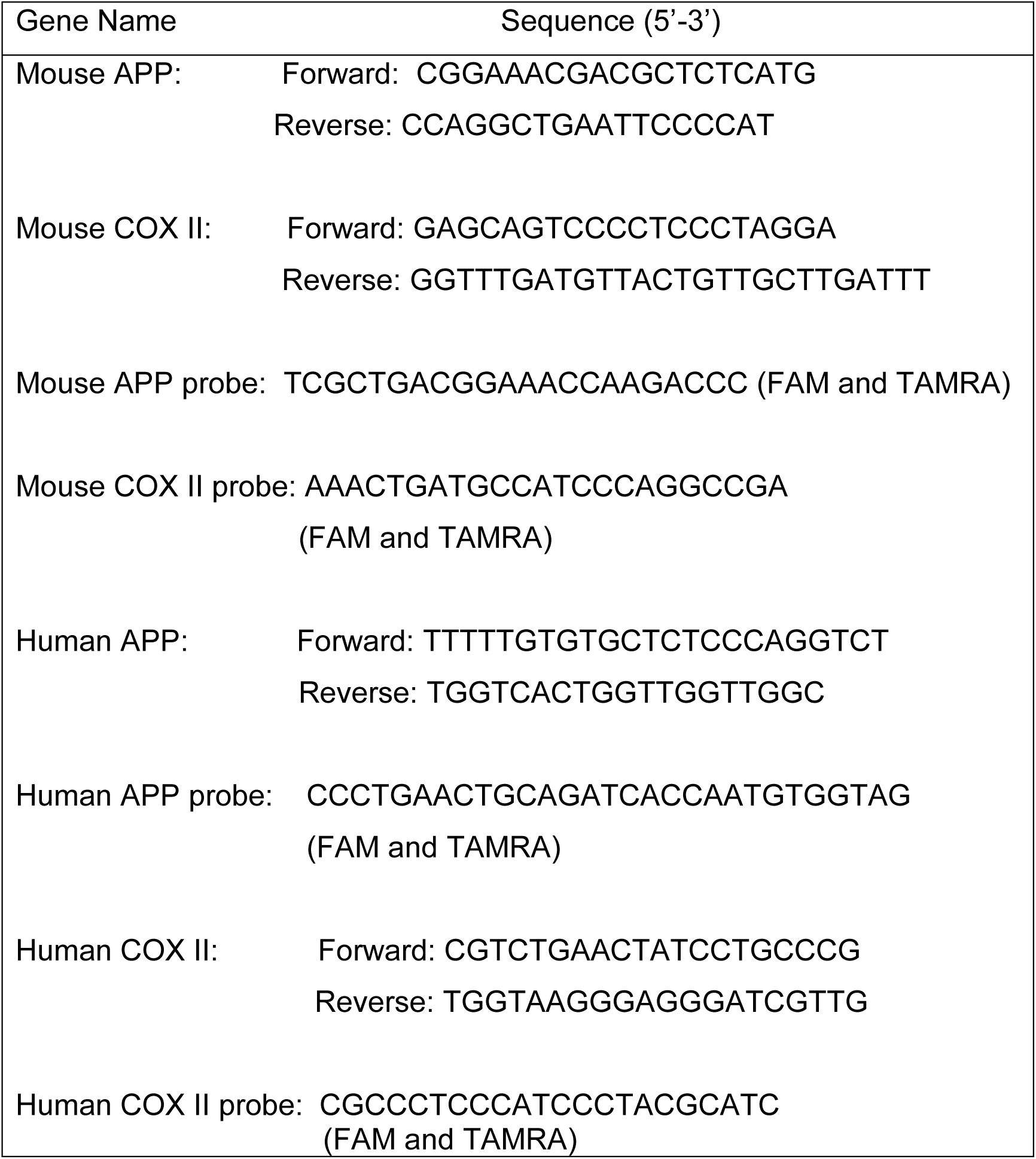
Primer Sequences.

### Imaging TNTs

Glass slides were coated with poly-L-lysine (10 μg/mL) for 2 hours before seeding freshly isolated monocytes (4,000/cm^2^). Monocytes were differentiated into M2-like Mφs for 10 days with 100 ng/mL M-CSF. Leukemic cells (KG-1a, THP-1, U937) were added at 10,000 cells per well and co-cultured for 48 hours. Cells were stained with 250 nM MTDR for 1 hour at 37°C, washed, and fixed with 4% v/v paraformaldehyde for 20 minutes. Permeabilisation was performed with 0.5% v/v Triton X-100 for 20 minutes, followed by staining with ActinGreen 488 and Hoechst 33342 (5 μg/mL) for 30 minutes. Vectashield with DAPI was applied. Imaging was performed on a Zeiss Axio Observer 7 using a 100x lens and images were captured with an Axiocam 705 mono camera on ZEN 3.5 software. Image analysis was conducted in Fiji. All procedures were carried out at room temperature unless otherwise specified, ensuring consistent staining quality and optimal imaging conditions throughout the experiment.

### Statistical analysis

All statistics tests are stated in the legends. A p-value of ≤0.05 was considered significant. All analysis was carried out using GraphPad Prism version 10 software (La Jolla, CA, USA).

## RESULTS

### M2-like Mφs protect AML cells from therapy-induced apoptosis

M2-like CD206^+^ Mφ precursor monocytes (17), and M2-like (CD163^+^CD206^+^) Mφs (6), are elevated in the BM of AML patients. We utilized M-CSF differentiated and polarized MDMs from healthy blood donors, as a model for M2-like Mφs (**Fig. 1A**). M-CSF polarized Mφs expressed CD206 to a similar extent as Mφs exposed to IL-4 and IL-13. Furthermore, M-CSF polarized Mφs expressed low levels of CD86, comparable to that of IL-4 and IL-13 polarized Mφs (**Supp. Fig. S1A**). The M2-like phenotype was further confirmed, by analyzing conditioned media from the M-CSF polarized Mφs for several chemokines and cytokines. M-CSF polarized Mφs secreted high levels of M2-associated chemokines (e.g. CCL22 and CCL24) (**Supp. Fig. S1B**), and chitinase-like protein 3 (CHI3L1) (**Supp. Fig. S1C**), consistent with reports that M-CSF differentiated Mφs resemble M2-like Mφs/tumour-associated macrophages (1, 18–20). We and others have shown that M2-like Mφs significantly contribute towards the development of treatment resistance in various haematological malignancies, including multiple myeloma (MM) (21–24), with M2-like Mφs shielding AML cell lines from apoptosis induced by targeted therapies (9, 10). However, the capacity of M2-like Mφs to protect AML cells, from DNR treatment is unknown. To test this, a layered Mφ-AML co-culture system was established (**Fig 1A**). Direct contact with M2-like Mφs significantly attenuated DNR-induced apoptosis in all AML cell lines investigated (**Fig. 1B-D**). Furthermore, Mφ-CM was used to ascertain if Mφ secreted factors exhibited chemoprotective properties. This protection was observed in two out of the three AML cell lines cultured with Mφ-CM.

**Fig. 1:**
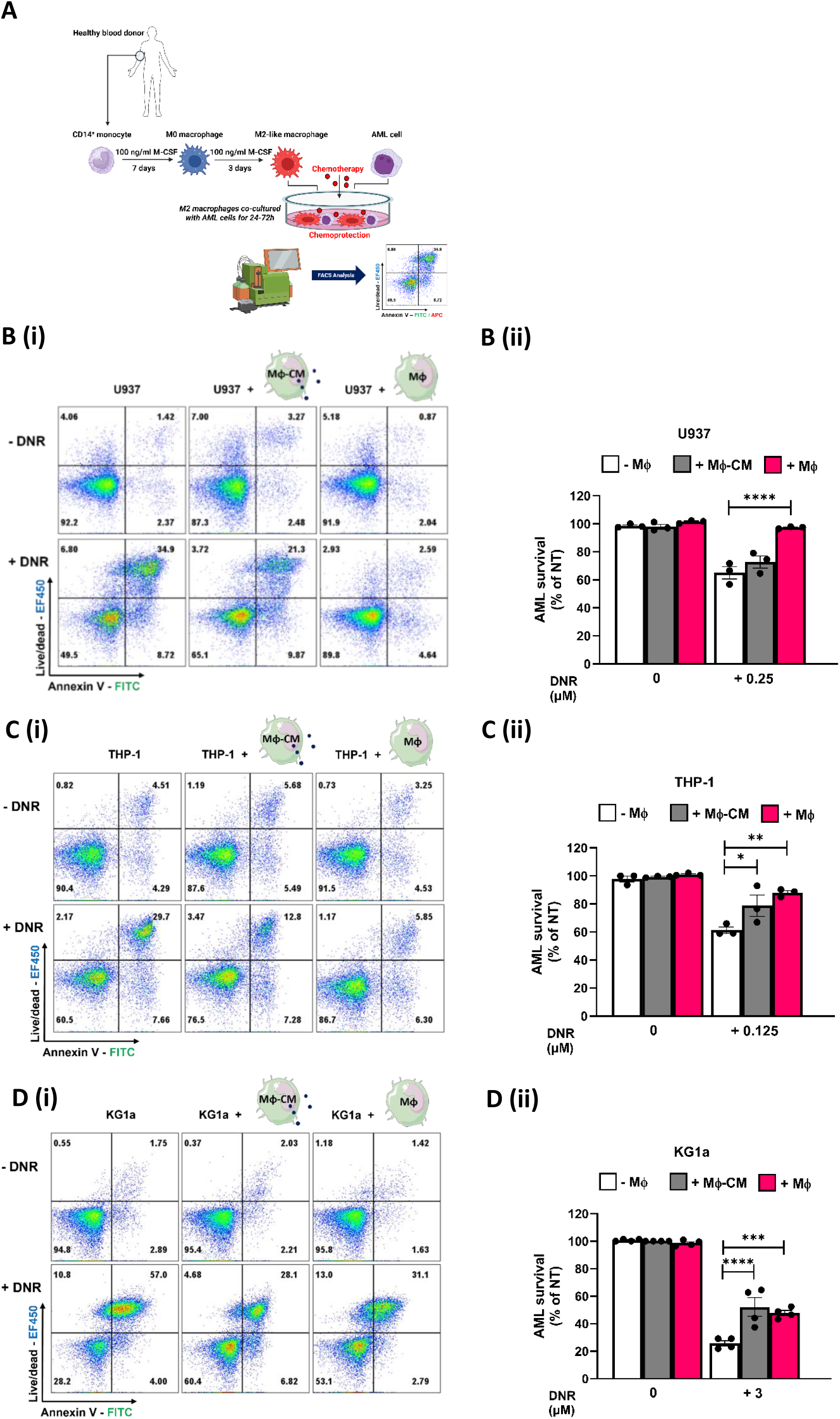
M2-like Mφs confer protection to AML cells from chemotherapy-induced apoptosis. (A) Diagram illustrating how CD14^+^ monocytes were differentiated to MDMs and polarized. CD14^+^ monocytes were isolated from healthy blood donors, as a model for M2-like Mφs. AML cells were treated with DNR either in monoculture (-Mφ), in the presence of 50% macrophage conditioned media (Mφ-CM) or in direct co-culture with macrophages (Mφ). Representative plots of flow cytometric analysis of Annexin V-FITC and Viability Dye eFluor™ 450 fluorescence are shown for each cell line (Bi, Ci and Di). The treatment conditions are as follows; (Bii) U937: 0.25μM DNR for 24 hours (n=3), (Cii) THP-1: 0.125μM DNR for 72 hours (n=3), (Dii) KG-1a: 3μM DNR for 48 hours (n=4). AML survival (% of non-treated [NT]) was determined by staining with Fixable Viability Dye eFluor™ 450 and Annexin V-FITC. Cells were analysed using the MACSQuant® Analyzer 10. Data are Mean ± SEM and were analysed using a one-way ANOVA followed by Tukey’s multiple-comparison test, *p<0.05, **p<0.01, ***p<0.001, ****p<0.0001.

Together, these findings demonstrate that M2-like Mφs protect a diverse range of AML subtypes from therapy-induced apoptosis mostly via cell-to-cell contact.

### AML therapies enhance mitochondrial transfer from M2-like Mφs to AML cells via TNTs with inhibition of TNTs suppressing Mφ-driven chemoprotection

Findings from the Schuringa research group demonstrate that M2d (IL-6 polarized) Mφs transfer functional mitochondria to primary AML cells under steady-state conditions, enhancing metabolism/ATP production and subsequently the proliferation of AML blasts (8). However, the effect of chemotherapeutic stress on mitochondrial transfer from M2-like macrophages (Mφs), the mechanisms by which M2-like Mφs transfer mitochondria to AML cells, and whether this transfer underpins Mφ-driven therapy resistance in AML remain unclear. We hypothesized that DNR and Ara-C enhance mitochondrial transfer from Mφs to AML cells and that this transfer mediates Mφ-driven chemoprotection, based on previous findings (11, 25). We first assessed whether M-CSF-polarized M2-like Mφs transferred mitochondria to AML cell lines and primary cells (**Fig. 2A**). MitoTracker studies indicated that, under steady-state conditions, mitochondria were transferred from M2-like Mφs to all AML cell lines examined, with transfer highest in THP-1, followed by U937 and KG1a (**Fig. 2B**). DNR alone and in combination with Ara-C further enhanced mitochondrial transfer from M2-like Mφs to AML cell lines (**Fig. 2C**) and primary AML cells (**Fig. 2D**). We then examined tunnelling nanotubes (TNTs), which are known to mediate mitochondrial transfer from donor to acceptor cells (26), including transfer from BM stromal cells to AML cells (27, 28).

**Fig. 2:**
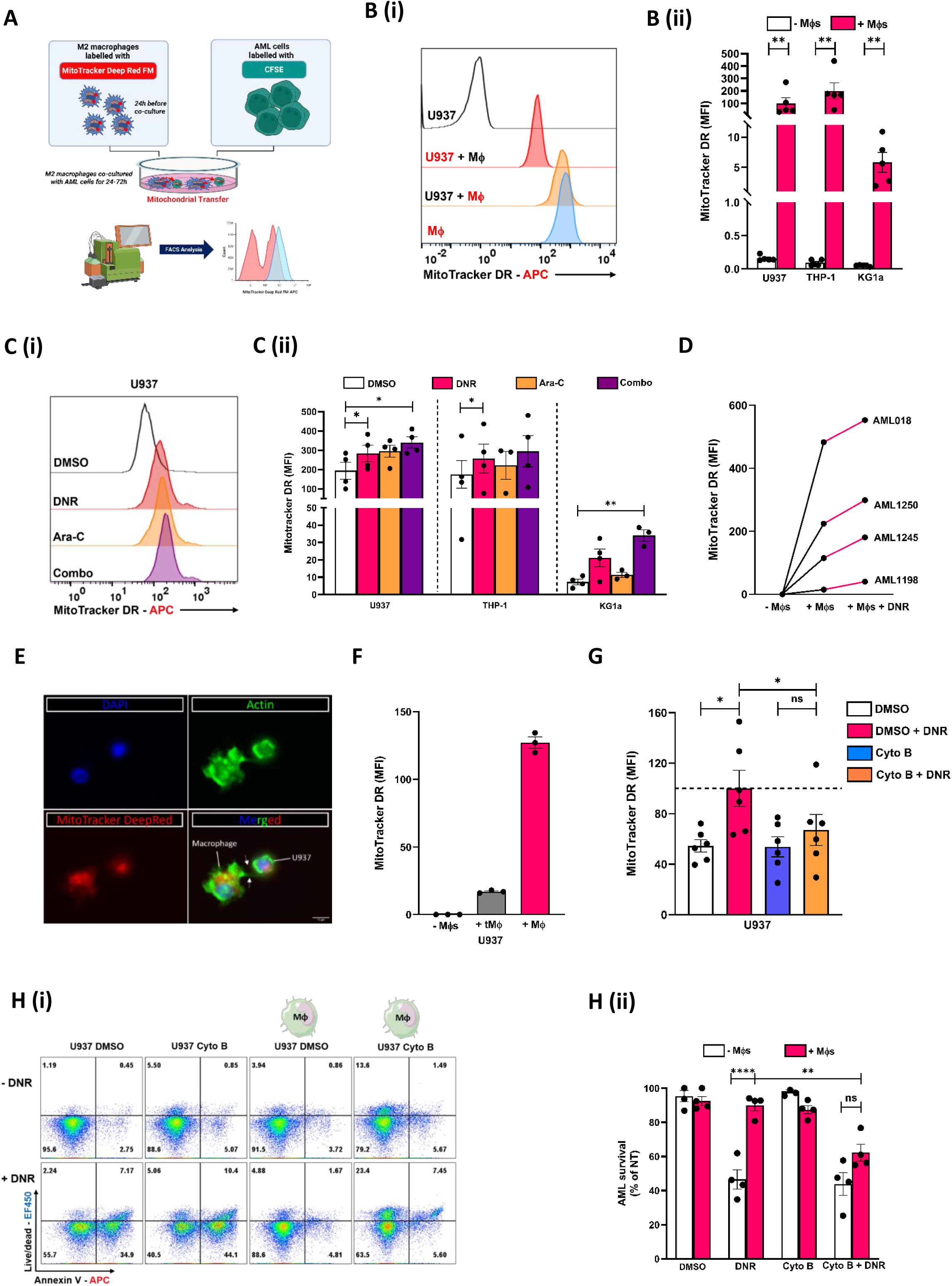
AML therapies enhance the transfer of mitochondria from M2-like monocyte-derived Mφs to AML cells, with mitochondrial transfer between macrophages and AML cells likely TNT-mediated. (A) Mφs were loaded with MTDR on Day 9 of culture of M2-like Mφs. CFSE-labelled U937, KG1a and THP-1 were then co-cultured with Mφs at a 1:1 ratio with M2-like Mφs for 24, 48 and 72 hours respectively. Mitochondrial transfer was then assessed in CFSE-labelled FVD^-^ viable AML cells, with MTDR analysed in the R1 (APC) channel, via the MACSQuant® Analyzer 10. (Bi) Representative histograms of flow cytometric analysis of MTDR fluorescence, U937 = unlabeled U937 cells; U937 + Mφ = MTDR signal from U937 cells in co-culture with M2-like Mφs; U937 + Mφ = MTDR signal from M2-like Mφs in co-culture with U937 cells and Mφ = MTDR signal from M2-like Mφs. (Bii) Data are Mean ± SEM of n=5 and analysed by a Mann-Whitney U Test, **p<0.01. (Ci) Representative histograms of flow cytometric analysis of MTDR fluorescence. (Cii) AML cell lines were cultured on and off MTDR loaded Mφs in the presence or absence of DMSO (vehicle control) or DNR or Ara-C or the combination of DNR & Ara-C, for the indicated times and concentrations: U937: 0.25μM DNR for 24 hours; THP-1: 0.125μM DNR for 72 hours; KG-1a: 3μM DNR for 48 hours, the same times for Ara-C at 2.5μM for all AML cell lines (n=3-4). Data were then analyzed by a one-way ANOVA followed by Dunnett’s multiple-comparison test, *p<0.05, **p<0.01. (D) Primary AML cells were cultured on and off MTDR loaded Mφs in the presence or absence of 0.125μM of DNR for 24h and mitochondrial transfer assessed as above (n=4). (E) U937 cells can interact with macrophages via TNTs. Blue (top left) is indicative of DNA staining (DAPI), green (top right) is the actin staining (ActinGreen488), red (bottom left) corresponds to mitochondria (MTDR), and the merged image is shown bottom right (n=1). Scale bar = 10 μm. White arrows indicate TNTs. (F) U937 cells were cultured alone or cultured either indirectly (tMφ, via transwell inserts) or directly with MTDR loaded Mφs for 24h. (G) U937 cells were cultured on MTDR loaded Mφs in the presence or absence of DNR or 1 μM Cyto B or the combination of DNR & Cyto B, for 24 hours. Mitochondrial transfer was then assessed in CFSE-labelled AML cells, with MTDR analysed in the R1 (APC) channel, via the MACSQuant® Analyzer 10 (n=6), and analysed by a one-way ANOVA followed by Tukey’s multiple-comparison test, *p<0.05, ns = non-significant. (Hi) Representative plots of flow cytometric analysis of Annexin V-FITC and Viability Dye eFluor™ 450 fluorescence. (Hii) U937 cells were treated with DNR +/- cyto B either in monoculture (-Mφ) or in direct co-culture with macrophages (+Mφ) for 24h. AML survival (% of NT) was determined by determined by staining with Fixable Viability Dye eFluor™ 450 and Annexin V-APC. Cells were analysed using the MACSQuant® Analyzer 10 (AML cell lines). (H) Data are Mean ± SEM of n=4 and analysed by a two-way ANOVA followed by followed by Tukey’s multiple comparisons test, **p<0.01, ***p<0.0001, ns = non-significant, ns = non-significant.

We predicted that M2-like Mφs transfer mitochondria via TNTs. TNTs exhibit dynamic structural and functional changes; thus, we used fixed-cell imaging to monitor transfer. M2-like Mφs were cocultured with leukemic cells for 48 hours, mitochondria in both cell types were labelled with MTDR. Fluorescent microscopy revealed TNT connections observed between U937 (**Fig. 2E**), THP-1, and KG1a cells and M2-like Mφs (**Supp. Fig. S2A, B**), with TNT connections formed between leukemic cells Mφs (**Supp. Fig. S2C**) and between Mφs (**Supp. Fig. S2D**). MTDR signal in AML cells was nearly abolished when AML cells were physically separated from Mφs using transwell inserts (**Fig. 2F**), supporting direct contact-dependent mitochondria transfer. Treatment with cytochalasin B, which blocks F-actin polymerization critical for TNTs(29), reduced DNR-enhanced mitochondrial transfer to U937 cells by ∼34% (**Fig. 2G**). Exposure of AML cells to conditioned media from MTDR-loaded Mφs produced low MTDR signals, potentially indicating minor transfer via contact-independent mitochondria transfer potentially via extracellular vesicles (EVs), which can mediate mitochondrial transport through EV-cell fusion (30). Finally, we generated iPSC-derived M2-like Mφs from the human skin fibroblast line SFCi55, previously used to produce terminally differentiated M1 and M2-like Mφs (14), providing an additional Mφ model to further study mitochondrial transfer and its role in therapy resistance in AML. To ensure the successful generation of monocyte/macrophage-like precursors (iMacs) from the SFCi55 cells, we assessed cell morphology and also marker expression via light microscopy and FACS, to confirm that the iMacs were CD45^+^, Podoplanin (PDPN^-^) and thus not fibroblasts, but immune cells differentiating along the haematopoietic lineage.

The iMac cells displayed morphological features consistent with monocytic cells (**Supp. Fig. S3A**), and were CD14^+^, CD115^+^, CD11b^+^, and CD64^+^, thus demonstrating successful differentiation along the myeloid lineage (**Supp. Fig. S3B**). As with donor-derived M2-like Mφs, M2-like iPSC-derived Mφs had the capacity to protect U937 cells from DNR-induced apoptosis (**Supp. Fig. S3C**). Using M2-like human iPSC-derived Mφs (**Supp. Fig. S3D**) and M2-like murine BMDMs (produced from C57BL/6 mice), we confirmed findings from our experiments using primary monocyte-derived M2-like Mφs (**Supp. Fig. S4**). Given that cytochalasin B suppressed DNR-enhanced mitochondrial transfer from Mφs to AML cells, we investigated the hypothesis that cytochalasin B exposure could abolish the ability of Mφs to protect AML cells from DNR induced killing. In fitting with this prediction, cytochalasin B was able to resensitise U937 cells to the killing effects of DNR when cultured with Mφs (**Fig. 2H**). Given that M2d Mφs donate mitochondria to primary AML cells, and that co-culture of primary AML cells with M2-like Mφs enhances AML cell proliferation (8), we assessed the proliferation of AML cells cultured alone or with M2-like Mφs. Culture of AML cells with M2-like Mφs did not impact AML cell proliferation (data not shown).

Collectively these data suggest that mitochondrial transfer from Mφs to AML cells is enhanced by conventional AML therapies, is dependent on Mφ-to-AML cell contact, and most likely occurs via TNTs.

### Mitochondria transferred to AML cells are functionally active contributing to increased metabolic capacity and reduced ROS generation

We next investigated the function of the transferred mitochondria. We found that following co-culture with M2-like primary monocyte-derived Mφs, AML cells exhibited increased basal and maximum mitochondrial respiration, as well as increased spare respiratory capacity and elevated ATP generation compared with monocultured AML cells (**Fig. 3A**). We then tested the hypothesis that AML cells co-cultured with Mφs would display lower ROS levels under steady-state and DNR-induced stress conditions. This was based on mitochondria possessing key anti-oxidant systems (e.g. superoxide dismutase [SOD]) (31), and that acquisition of chemoresistance has been reported to correlate with reduced ROS levels (32). Assessment of total ROS levels showed that U937 cells cultured with M2-like Mφs exhibited reduced ROS generation under both steady-state and DNR-driven conditions, as compared to U937 monocultures (**Fig. 3B**).

**Fig. 3:**
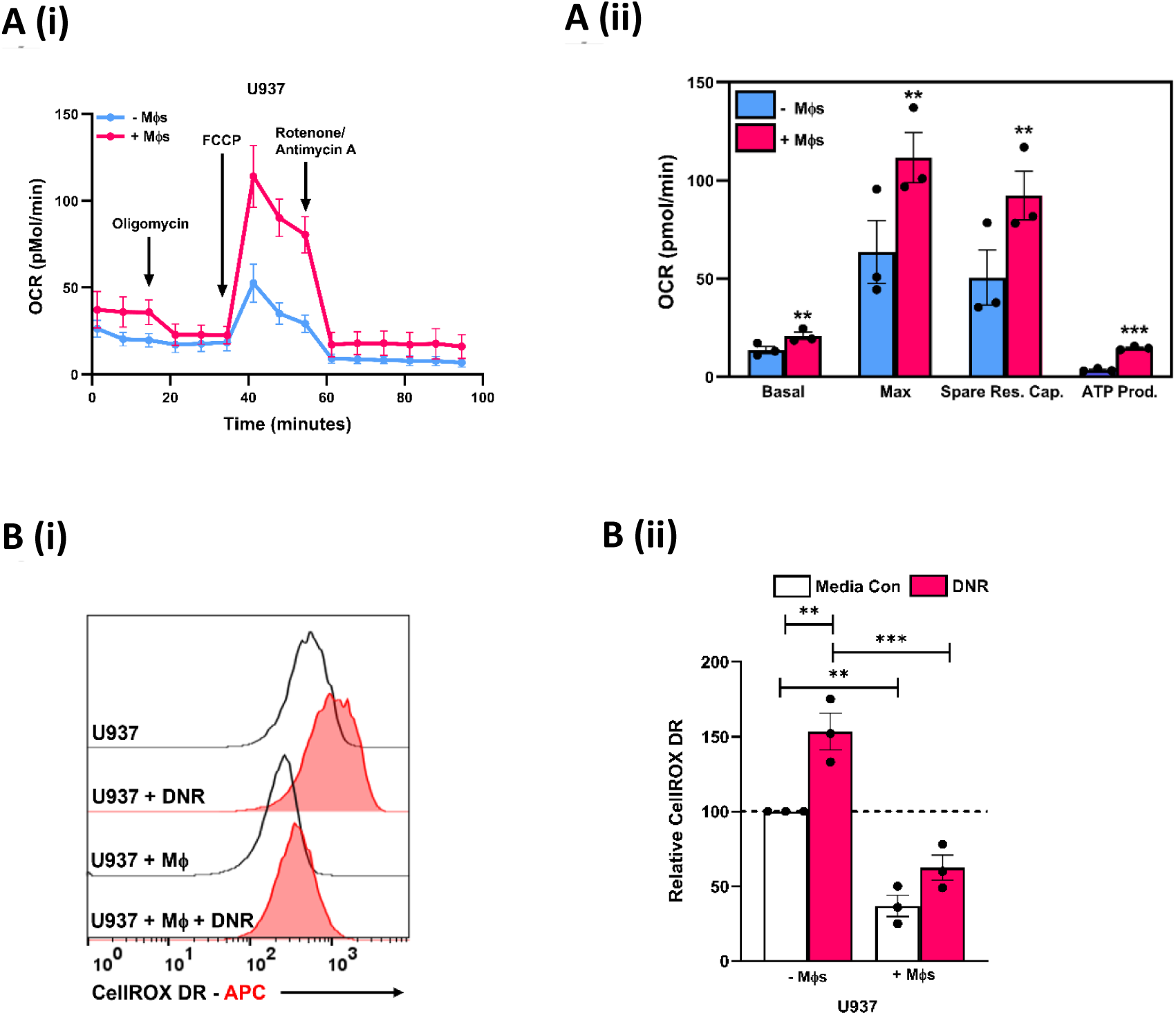
Mitochondria hijacked by AML cells are functionally active contributing to increased metabolic capacity and reduced ROS generation. (Ai) U937 cells were grown with and without M2-like monocyte-derived Mφs for 24 hours and then analyzed independently, using the Seahorse XFp Analyzer with the Mito Stress Test Kit. Sequential injections of Oligomycin (O), carbonyl cyanide-4-(trifluoromethoxy)phenylhydrazone (F), and Rotenone (R) were used to obtain respiration dynamics presented in panel (Aii). Data are mean ± SEM of n=3 and were then analyzed by using a paired student’s t-test, **p<0.01. (Bi) Representative histograms of flow cytometric analysis of CellROX fluorescence. (Bii) CFSE-labelled U937 were co-cultured at a 1:1 ratio with M2-like Mφs and in the presence of absence of DNR for 24 hours. Total ROS levels were then assessed in CFSE-labelled U937 cells using CellROX, which was analysed in the R2 (APC-Cy7) channel, via the MACSQuant® Analyzer 10. Relative CellROX values were calculated as a percentage of values obtained for U937 cells cultured alone in cRPMI (media con). The data are Mean ± SEM of n=3 and analysed by a two-way ANOVA followed by Tukey’s multiple-comparison test, **p<0.01, **p<0.001.

Taken together, mitochondria acquired by AML cells from M2-like Mφs are functional, contribute to the energy requirements of AML cells, and reduce ROS levels.

### LIMK inhibition and Metformin block daunorubicin-induced mitochondrial transfer and Mφ-driven therapy resistance

To identify proteins involved in Mφ-driven mitochondrial transfer and therapy resistance in AML, U937 cells were cultured alone or with M2-like Mφs for 24 hours. AML cells were then isolated and analysed using TMT-MS proteomics.

Five proteins, SNRPD3, COLGALT1, PDCD6IP, RHOC, and H2AX were upregulated in AML cells following co-culture (**Fig. 4A**). We next examined, via gene expression profiling interactive analysis (GEPIA), whether genes that encode these proteins correlated with clinical outcomes in AML using TCGA data. Genes that encode RHOC and SNRPD3 proteins were associated with reduced OS (**Fig. 5B & Supp. Fig. S5**). Considering the RhoC-LIMK-cofilin pathway’s role in TNT formation and mitochondrial transfer (33), and that genotoxic stress such as doxorubicin can activate this pathway (34), we hypothesized that DNR triggers RhoC-LIMK signalling. This leads to phosphorylation and inactivation of the F-actin depolymerizing protein cofilin by LIMK1/2, promoting TNT formation and mitochondrial transfer. Elevated cofilin expression also correlated with inferior overall survival (**Fig. 4C**). DNR induced cofilin phosphorylation, peaked at 12 hours post-treatment (**Fig. 4D**). LIMK2 mRNA was significantly higher in AML samples than in normal mononuclear cells (GEPIA) (**Fig. 4E**). To test if DNR-induced cofilin phosphorylation occurs via LIMK1/2, we used the selective inhibitor TH-257 (35). LIMK inhibition reduced DNR-induced cofilin phosphorylation (**Fig. 4F**), suppressed DNR-enhanced mitochondrial transfer from Mφs to AML cells (**Fig. 4G**), and diminished Mφ-mediated chemoprotection (**Fig. 4H**). Importantly, TH-257/LIMK inhibition did not affect the survival of M2-like Mφs (**Supp. Fig. S6A**). Given that TH-257 is a tool compound, and that no LIMK inhibitors to date have entered the clinic, we reviewed the literature to ascertain FDA approved drugs that have been shown to block mitochondrial transfer between stromal cells and AML cells for the potential for drug repurposing.

**Fig. 4:**
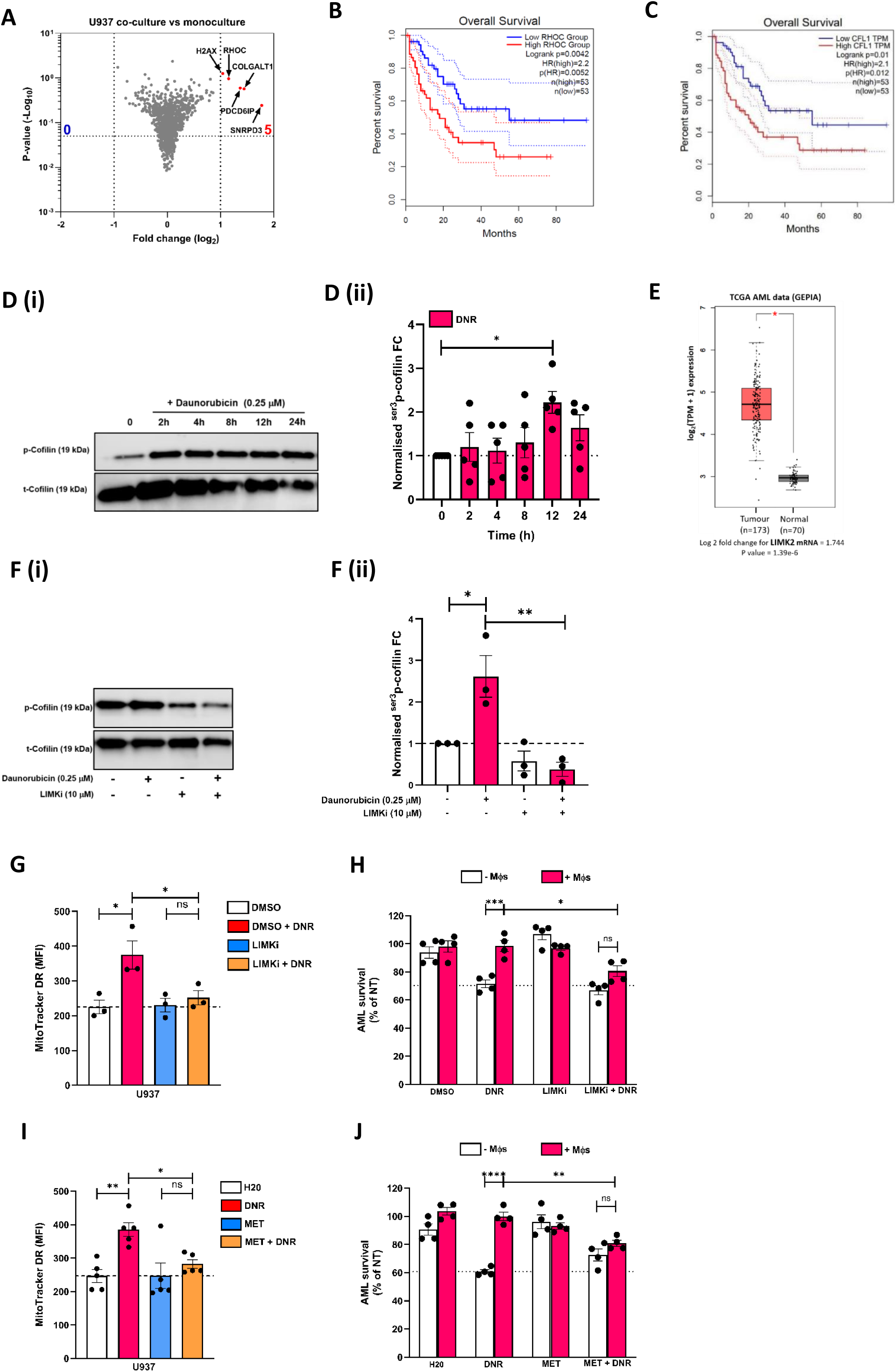
LIMK inhibition and Metformin suppress daunorubicin-induced mitochondrial transfer and Mφ-driven therapy resistance. (A) CFSE-labelled U937 cells were directly cultured either in the presence (co-culture) or absence (monoculture) of M2-like Mφs for 24h. CD14^-^ U937 AML cells were then isolated from CD14^+^ Mφs, via the use of CD14 magnetic activated cell sorting. Proteomic profiling was then conducted on purified AML whole cell lysates (WCLs), via a TMT11-plex. Data shows the average values of n=4. (B) Association of transcript expression levels of RhoC and (C) Cofilin-1 with overall survival of patients with AML, plots were generated from interrogation of TCGA data via the GEPIA database. (Di) U937 were exposed to 0.25 mM DNR for up to 24h. Levels of ^ser3^p-cofilin and total cofilin (t-cofilin) were then assessed in purified U937 WCLs by immunoblotting. (Dii) Fold change of normalised ^ser3^p-cofilin to t-cofilin densitometry values are shown. The data are Mean ± SEM of n=5 and analysed by a one-way ANOVA followed by Dunnett’s multiple comparisons test, *p<0.05. (E) TCGA data were analyzed using GEPIA, and the expression of LIMK2 in AML compared to the normal samples is shown. (Fi) U937 were exposed to 0.25 mM DNR in the presence or absence of DMSO or 10mM of TH-257 for 12h. Levels of ^ser3^p-cofilin and total cofilin (t-cofilin) were then assessed in purified U937 WCLs by immunoblotting. (Fii) Fold change of normalised ^ser3^p-cofilin to t-cofilin densitometry values are shown. The data are mean ± SEM of n=3 and analysed by a one-way ANOVA followed by a Tukey’s multiple comparisons test, *p<0.05, **p<0.01. (G) CFSE-labelled U937 cells were cultured in the presence or absence of DNR, DMSO or 10 mM TH-257 either off (-Mφ) or on (+Mφ) MTDR loaded Mφs in the presence or absence of DNR for 24 hours. Mitochondrial transfer was then assessed in CFSE-labelled AML cells, with MTDR analysed in the R1 (APC) channel, via the MACSQuant® Analyzer 10 (n=3), and analysed by a one-way ANOVA followed by Tukey’s multiple comparisons test, *p<0.05, ns = non-significant. (H) AML survival (% of NT) was determined by staining with Fixable Viability Dye eFluor™ 450 and Annexin V-APC. Data are Mean ± SEM of n=4 and were then analyzed using a two-way ANOVA followed by Tukey’s multiple comparisons test, *p<0.05, ***p<0.001, ns = non-significant. (I) CFSE-labelled U937 cells were cultured in the presence or absence of DNR, water or 5 mM metformin either off (-Mφ) or on (+Mφ) MTDR loaded Mφs in the presence or absence of DNR for 24 hours. Mitochondrial transfer was then assessed in CFSE-labelled AML cells, with MTDR analysed in the R1 (APC) channel, via the MACSQuant® Analyzer 10 (n=5) with the data analysed by a one-way ANOVA followed by Tukey’s multiple comparisons test, *p<0.05, **p<0.01, ns = non-significant. (J) AML survival (% of NT) was determined by staining with Fixable Viability Dye eFluor™ 450 and Annexin V-APC. Data are Mean ± SEM of n=4 and were then analyzed using a two-way ANOVA followed by Tukey’s multiple-comparison test, **p<0.01, p<0.0001, ns = non-significant.

We identified metformin, which inhibits the transfer of mitochondria from bone marrow stromal cells (BMSCs) to AML cells, and overcomes BMSC-driven chemoresistance in AML cells (36). Given that Metformin has been shown to reduce thrombin-induced p-cofilin levels, via an increase in PP2A activity (37), we assessed the impact of metformin on steady-state and DNR-induced p-cofilin levels. Metformin did not decrease steady-state or DNR-induced ^ser3^p-Cofilin levels (data not shown). However, metformin did reduce mitochondrial transfer from Mφs to AML cells (**Fig. 4I**), and resensitized AML cells to the killing effects of DNR when cultured with Mφs (**Fig. 4J**). Moreover, metformin did not affect the survival of M2-like Mφs (**Supp. Fig. S6B**).

Together these findings demonstrate that DNR-enhanced mitochondrial transfer and Mφ-driven chemoprotection is partly dependent on the LIMK-Cofilin pathway.

## DISCUSSION

Here we sought to ascertain whether M2-like macrophages protect AML cells from apoptosis induced by standard therapies, whether this resistance depends on the LIMK-Cofilin axis, and if targeting mitochondrial transfer or LIMK-Cofilin could be potentially therapeutic. CD206⁺ Mφs protected AML cells from DNR primarily via direct cell contact. This aligns with reports showing M2-like Mφs confer resistance to venetoclax and midostaurin, which is lost when Mφs are separated from AML cells (10). Consistent with prior studies (8), we found M2-like Mφs donate mitochondria to AML cells, which we confirmed using iPSC-derived Mφs and murine BMDMs. DNR alone and in combination with Ara-C combination enhanced mitochondrial transfer from M2-like Mφs to AML cells. This corroborates previous reports showing that DNR enhanced mitochondrial transfer from murine and human BMSCs to AML cells respectively (11, 25). Furthermore, we showed that mitochondrial transfer was largely dependent on Mφ-to-AML cell contact. In addition, transfer likely occurred via the formation of TNTs, as evidenced by the TNT-like structures observed when M2-like Mφs interact with AML cells, with reduced mitochondrial transfer occurring when the Mφs and AML cells were separated by transwell inserts or via the use of cyto B respectively. These findings agree with reports that mitochondrial transfer from BMSCs to AML cells is TNT-dependent (25). Our observations are therefore consistent with distinguishing features of TNTs (38). Our findings indicate that Mφ-driven therapy resistance is mediated by mitochondrial transfer, as cytochalasin B resensitized U937 cells to DNR, consistent with studies in MCF7 breast cancer cells acquiring chemoresistance through mitochondrial uptake (39).

We found that mitochondria transferred from M2-like Mφs to AML cells are functionally active, aligning with prior reports (8, 25). Additionally, M2-like Mφs reduced DNR-induced ROS in AML cells, consistent with observations in ALL where stromal mitochondrial transfer lowers ROS and protects blasts from chemotherapy-induced apoptosis (40). We identified upregulation of SNRPD3 and RHOC in AML cells following co-culture with M2-like Mφs, with high expression associated with poor survival. We further showed that DNR activates the RhoC-LIMK-cofilin pathway, and that LIMK inhibition or metformin reduces DNR-enhanced mitochondrial transfer, overcoming Mφ-mediated chemoprotection. Notably, FLT3-ITD AML patients taking metformin showed improved survival, suggesting broader therapeutic benefits (41). It is tempting to speculate that the benefits of metformin may extend to patients with other subtypes of AML, via the ability of metformin to block mitochondrial transfer between Mφs and AML cells.

Overall, our findings highlight Mφ-mediated mitochondrial transfer as a key driver of therapy resistance in AML and suggest targeting the RhoC-LIMK-Cofilin axis or mitochondrial transfer as promising therapeutic strategies.

## ACKNOWLEDGEMENTS

We thank Dr Kayleigh Thirlwell and Dr Kenneth Pallas for their assistance with Luminex analysis. We also thank Monica L. Guzman and Leonardo M. Martinez (Weill Cornell Medicine) for fruitful discussions regarding the project. We also thank Dr Richard Burchmore and Suzanne McGill (Glasgow Polyomics, University of Glasgow) for their help with analysis of the TMT-MS/proteomics data.

## AUTHOR CONTRIBUTIONS

CRediT (Contributor Roles Taxonomy) author statement: **EN, KM, AP, SH, KL, TF, KL, XB, RB :** Methodology, Investigation, Formal Analysis, Writing- Original Draft Preparation; **MD, AP, TH, and LM:** Methodology, Investigation, Formal Analysis, Writing-Review & Editing; **EN, KM, AP, CSG, LMF, KM, HW, KL, VAF, YK, MTSW:** Investigation, Writing- Review & Editing; **MTSW:** Methodology, Conceptualization, Investigation, Formal Analysis, Writing-Original Draft, Supervision, Project Administration, Funding Acquisition.

## FUNDING

MTSW was supported by Tenovus Scotland [S17-14 & S20-04], The British Society for Haematology [34847/ESR2017], The Sylvia Aitken Charitable Trust, The Miss AM Pilkington Charitable Trust and the Academy of Medical Sciences (AMS) Springboard scheme [SBF007\100123], which is joint-funded by the AMS, the Wellcome Trust, the Government Department of Business, Energy and Industrial Strategy (BEIS), the British Heart Foundation and Diabetes UK. TH is supported by the United Kingdom Medical Research Council [MR/X018512/1].

## DATA AND MATERIALS AVAILABILITY

All data required to assess the conclusions in the paper are present either within the paper itself or the included Supplementary Materials. Underlying raw data from this study are available from the corresponding author upon reasonable request.

## Notes

### Competing Interest Statement

The authors have declared no competing interest.

