## Supplemental figures for "LIMK Inhibition and Metformin Block Mitochondrial Transfer Overcoming Macrophage Driven Therapy Resistance in Acute Myeloid Leukaemia"

Nwarunma et al.

**Supp. Table 1:**

| Patient ID | Gender | Age | Karotype | Mutations |
|---|---|---|---|---|
| AML0018 | M | 68 | Normal | *NRAS, TET2, ASXL1 & EZH2* |
| AML1198 | F | 62 | Normal | *NRAS, GATA2, NPM1, DNMTA3A & IDH2* |
| AML1245 | F | 37 | 46,XX,del(7)(q22q32),t(8;21)(q22;q22),inv(18)(p11q21)[10] | *NRAS, BCORL1, IDH1* |
| AML1250 | F | 60 | 47,XX,+11[30] | *IDH2* |

### Supp. Figure S1

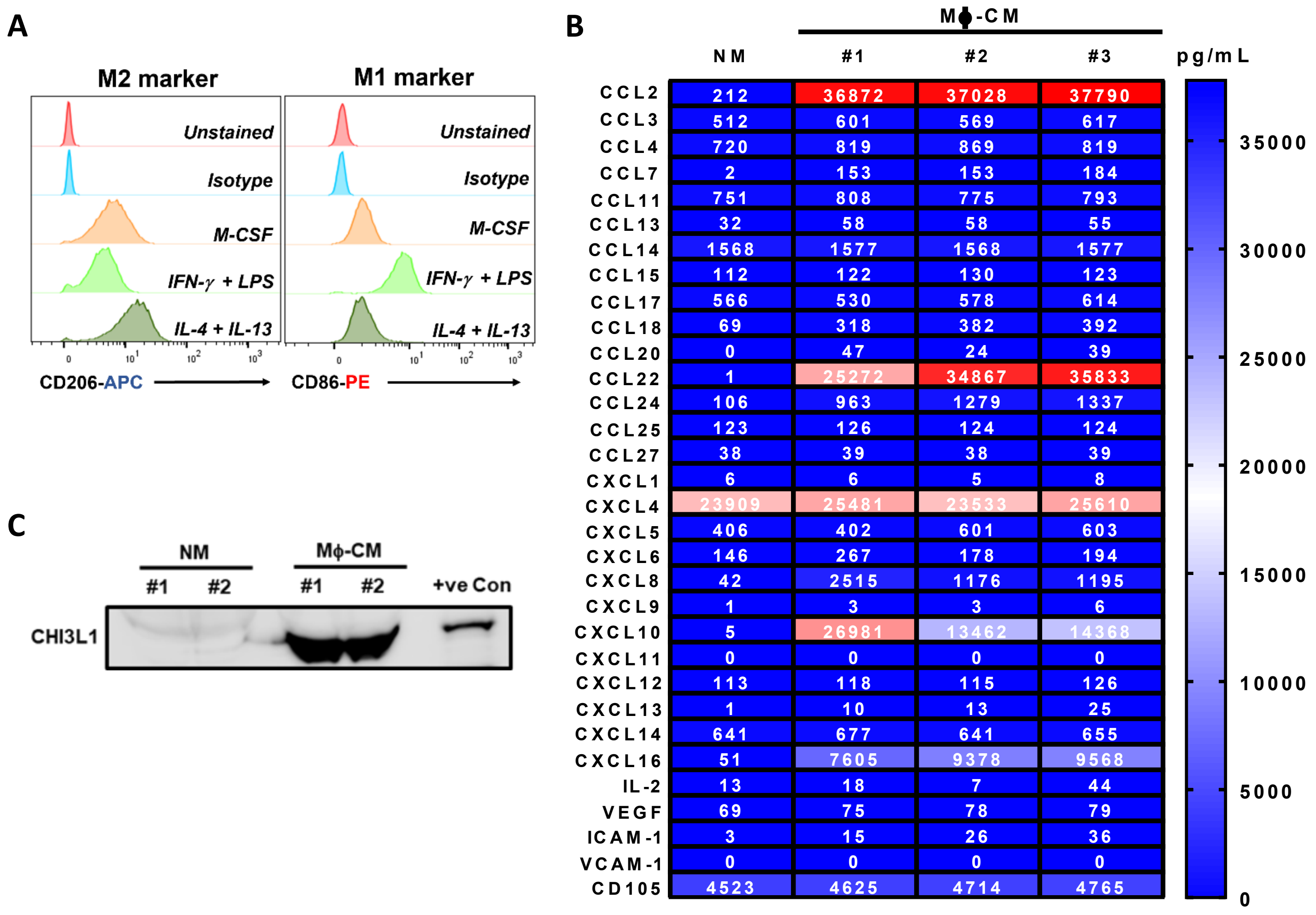


**Supplemental Figure S1: M-CSF differentiated and polarized MDMs resemble M2-like Mφs.** (A) Human CD14+ monocytes were differentiated into mature Mφs by culturing CD14+ monocytes with M-CSF (100 ng/mL) for 7 days. Mφs were then cultured for a further 24 hours with either M-CSF, or standard M1 (IFN-γ: 20 ng/mL & LPS: 100 ng/mL) and M2-(IL-4 & IL-13: 20 ng/mL each) polarising agents. Flow cytometry for M1 (CD86) and M2 (CD206) associated surface markers was performed. (B) Mature Mφs were exposed to 100 ng/mL of M-CSF for 3 days, following this cell culture supernatants were then harvested from the M2-like Mφs (Mφ-CM). Normal media (NM) and Mφ-CM were analyzed via a cytokine 30-Plex panel on a Luminex LX 200 Instrument. Concentrations determined from 1 batch of NM (n=1), and cell culture supernatants harvested from 3 healthy blood donors (n=3). (C) Normal media (NM) and Mφ-CM from 2 buffy coat donors (n=2), Mφ whole cell lysate (positive control = +ve con) were analysed for chitinase-like protein 3 (CHI3L1) via immunoblotting.

**Supp. Figure S2:**

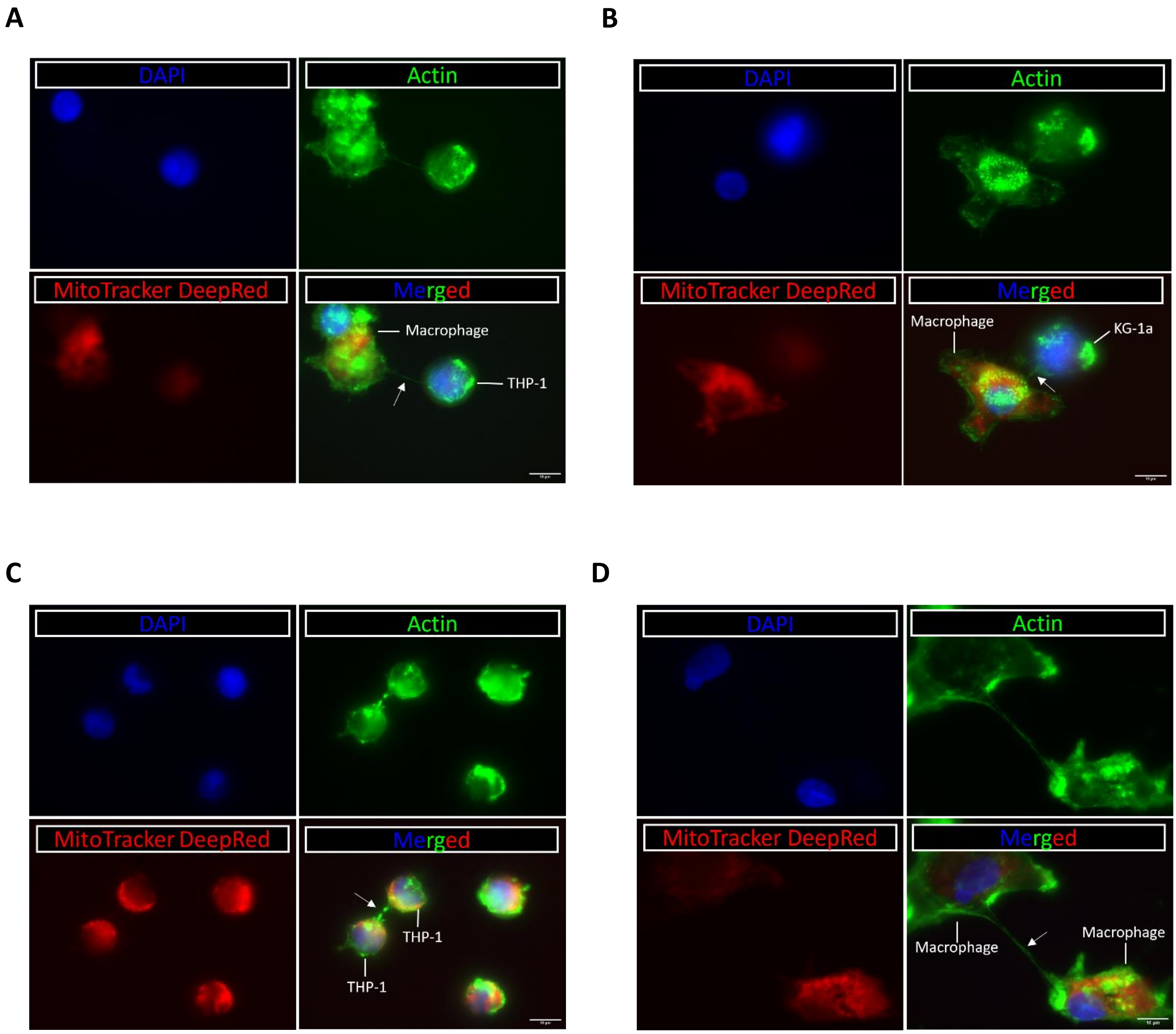


**Supplemental Figure 2: TNTs form between macrophages and AML cells, macrophages and leukaemic cells. (A)** THP-1 and **(B)** KG1a cells can interact with macrophages via TNTs. **(C)** THP-1 cells interact with each other, and **(D)** macrophages interact with each other via TNT formation. Blue (top left) is indicative of DNA staining (DAPI), green (top right) is the actin staining (ActinGreen488) and red (bottom left) corresponds to mitochondria (MitoTracker DeepRed). The merged image is shown bottom right. Scale bar = 10 μm. White arrows indicate TNTs.

**Supp. Figure S3:**

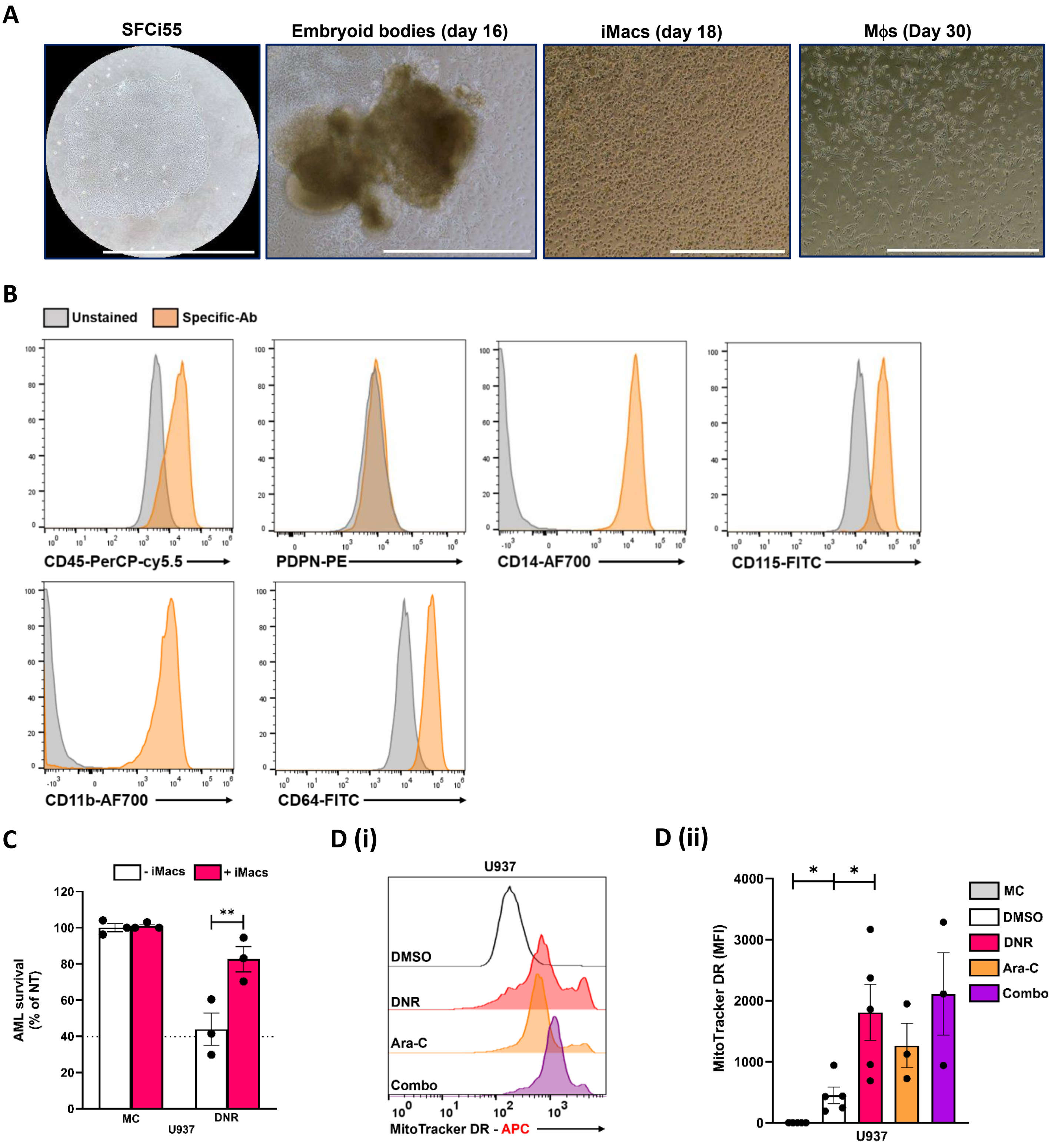


**Supplemental Figure 3: M2-like iPSC-derived Mφs transfer mitochondria to AML cells and protect AML cells from chemotherapy-induced apoptosis.** (A) Bright field images of human iPSC-derived macrophages taken at different stage of their differentiation/reprogramming into macrophages. Scale bar represents 5 μm. (B) Cell-surface marker expression (lineage and myeloid markers) in monocyte/macrophage-like precursors (iMacs) derived from SFCi55 iPSC, as assessed by flow cytometry. (C) CFSE labelled U937 cells lines were treated with 0.25 μM DNR either in monoculture (-iMacs) or in direct co-culture with iPSC-derived M2-like macrophages (+iMacs) for 24h.

AML survival (% of NT) was determined by determined by staining with Fixable Viability Dye eFluor™ 450 and Annexin V-APC. Cells were analysed using the MACSQuant® Analyzer 10. Data are Mean ± SEM of n=3 and were analysed by a two-way ANOVA followed by a Tukey's multiple comparisons test,**$p<0.01$. (D) U937 AML cells were cultured on and off (MC = monoculture) MitoTracker Deep Red FM loaded iPSC-derived M2-like Mφs in the presence or absence of 0.25μM DNR, or 2.5μM Ara-C or the combination of DNR & Ara-C for 24 hours (n=3-5). Mitochondrial transfer was then assessed in CFSE-labelled AML cells, with MitoTracker analysed in the R1 (APC) channel, via the MACSQuant 10 analyser, data were analysed by a one-way ANOVA followed by a Dunnett's multiple comparisons test, *$p<0.05$.

**Supp. Figure S4:**

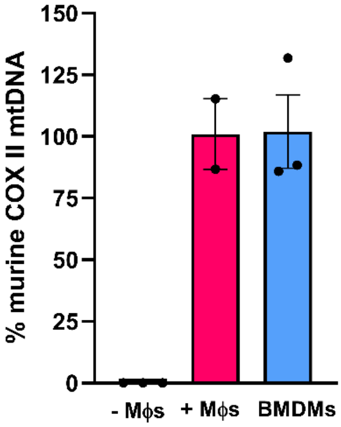


**Supplemental Figure 4: Murine bone marrow derived macrophages transfer mitochondria to AML cells.** U937 AML cells were cultured on and off M2-like murine BMDMs for 24 hours (n=2-3). The percentage of murine mitochondrial complex II was calculated by taking the negative log to the base 2 of the ΔΔCT of the monoculture and the coculture samples normalised to the human and murine nuclear DNA (APP).

**Supp. Figure S5:**

**A**

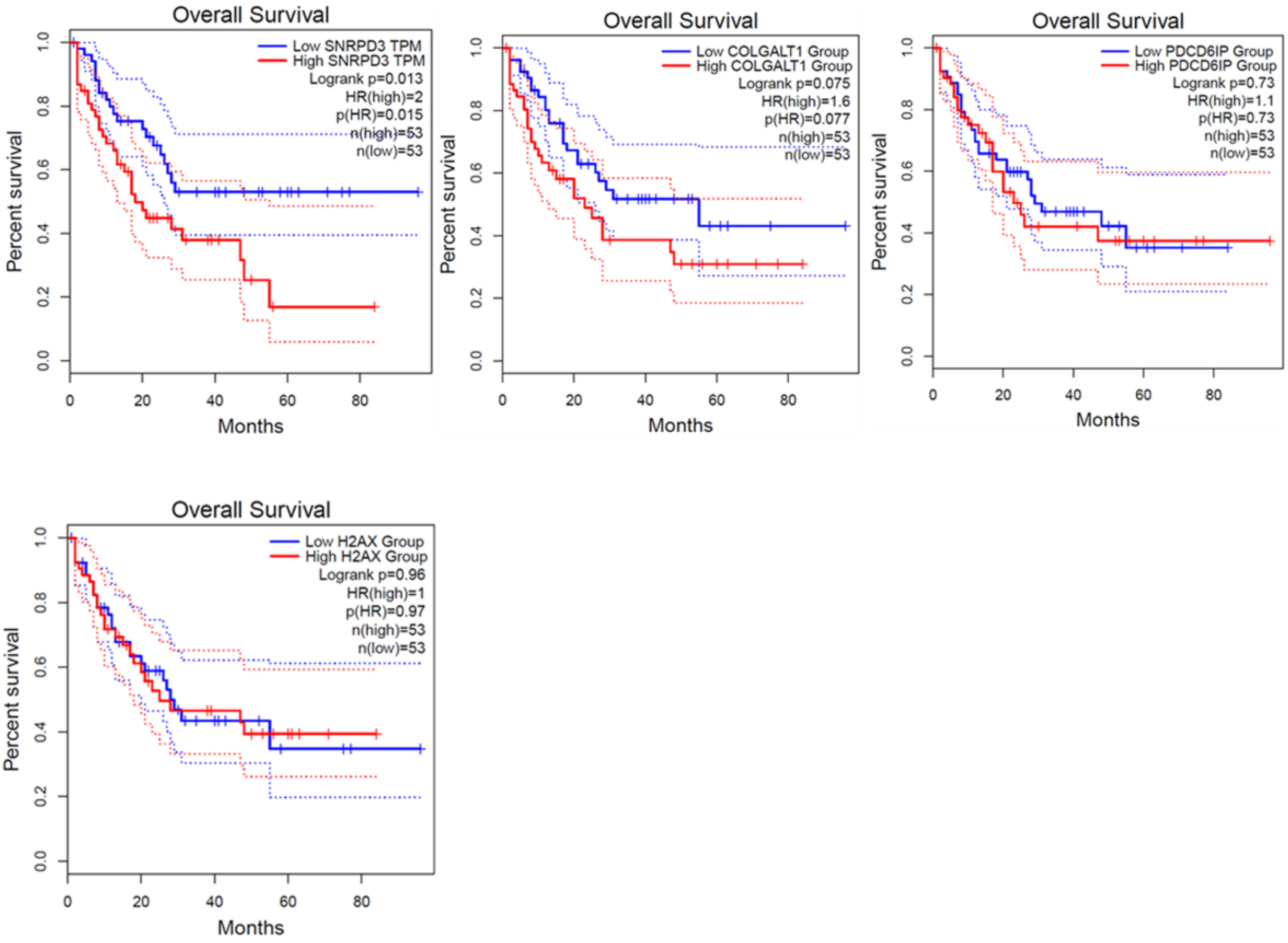


**Supplemental Figure 5: Correlation of transcript expression levels of protein "hits" with overall survival of AML patients. (A)** Data were obtained from the GEPIA database via interrogation of TCGA data.

**Supp. Figure S6:**

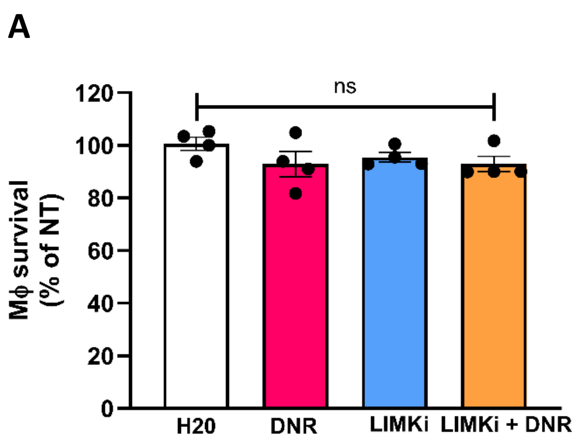


**Supplemental Figure 6: TH-257 and metformin do not impact the survival of M2-like Mφs.** CFSE-labelled U937 cells were cultured in the presence or absence of DNR, DMSO or water, 10 μM TH-257 or 5 mM metformin either off or on M2-like Mφs for 24 hours. Survival (% of NT) of $CFSE^{-}$ M2-like Mφs within the co-culture was determined by staining with Fixable Viability Dye eFluor™ 450 and Annexin V-APC. Data are Mean ± SEM of n=4 and n=5 and were analyzed by a one-way ANOVA followed by Dunnett's multiple-comparison test, ns= non-significant.
